# Effort in Oculomotor Control: Role of Instructions and Reward on Spatio-temporal Eye Movement Dynamics

**DOI:** 10.1101/2024.09.09.611577

**Authors:** Christian Wolf, Michael B. Steinborn, Lynn Huestegge

## Abstract

Effort is an important construct in several psychological disciplines, yet there is little consensus on how it manifests in behavior. Here, we focus on effort in terms of performance improvements beyond speed-accuracy tradeoffs and argue that oculomotor kinematics are conducive to a more fine-grained understanding of effortful behavior. Specifically, we investigated the efficiency and persistence of mere task instructions to induce transient effort. In a saccadic selection task, participants were instructed to look at targets as quickly and accurately as possible (standard instructions) or to mobilize all resources and respond even faster and more accurately (“*to give 110%*”, effort instructions). We compared results to standard speeded performance (baseline block) and to a potential upper performance limit linking effort to performance-contingent rewards (reward block). Performance improved beyond speed-accuracy tradeoffs when reward was present. Effort instructions reduced saccade latencies, increased amplitudes and changed the saccade main sequence relationship. Yet, these effects were more strongly pronounced and more persistent over time when effort was additionally rewarded. Altogether, the present findings underscore the possibility to intentionally activate cognitive resources for regulating oculomotor performance. Yet, its effectiveness and maintenance over time are more successful when behavior is rendered purposeful by the presence of reward.

**Significance Statement:** This study shows that people can exert voluntary control over the cognitive resources they devote to a task and that better task performance also goes along with changes in eye movement behavior. These findings thus suggest that eye movements are a particularly valuable tool when studying effort and cognitive control. Furthermore, eye movements can help to discriminate between phases in which people engage in a task and when they do not.

## Introduction

We complete some activities with ease, like watching a movie, whereas other tasks, like performing mental arithmetic, require mental effort. This particularly applies to tasks involving a high level of cognitive control. Performing these tasks for a longer duration can result in degraded performance (more errors, slower processing) and is possibly accompanied by the feeling of fatigue (Kok, 2022). Maintaining a desired level of performance can be achieved by taking a rest (Steinborn & Huestegge, 2016), using substances like caffeine (Connell et al., 2016), or by intentionally reinstating mental effort. Here, we investigated (i) whether and how long participants can intentionally restore effort merely based on instructions, (ii) how restored effort is modulated by the presence of additional reward, and (iii) how exactly these effects of effort are reflected in oculomotor markers, a response domain allowing for a particularly fine-grained analysis of a dynamic array of spatial and temporal aspects of performance.

### What is effort?

Effort is a prominent construct in a variety of disciplines, covering cognitive psychology, neuroscience, and behavioral economics (Verguts et al., 2015; Kurzban, 2016; Westbrook & Braver, 2015; 2016; Shenhav et al., 2017; Inzlicht et al., 2018; Silvetti et al., 2023). Although most laypeople have an intuitive understanding of effort, the concept is difficult to define from a scientific perspective, resulting in vague or even contradictory definitions (for an overview see Thomson & Oppenheimer, 2022; Shepherd, 2023). Yet, there is converging evidence for a few effort-related conceptualizations or findings that may lay the foundation for a minimal consensus.

First, mental effort can be considered aversive. Aversive signals in cognitive control have been suggested to mediate the invested effort or to motivate future task selection (Botvinick, 2007; Dreisbach & Fischer, 2011). In line with this, people typically prefer the task requiring less effort when confronted with a recurring choice between different tasks (Kool et al., 2010; McGuire & Botvinick, 2010; Dunn et al., 2016).

Second, incentives may motivate effortful behavior. Specifically, performance-contingent reward can simultaneously increase response accuracy and response speed in tasks requiring cognitive control (Krebs et al., 2010; Manohar et al., 2015; Wolf & Lappe, 2023), and rewarding effort can diminish or even overcome the avoidance of effortful tasks (Clay et al., 2022). Based on such findings, effort production is typically explained via a cost-benefit analysis (for an overview see Inzlicht et al., 2018; Otto & Daw, 2019; Székely & Michael, 2021): Tasks containing a response conflict (e.g. the Stroop task) require cognitive control, rendering them inherently costly. But if the benefits outweigh the costs, participants select more effortful tasks and/or activate cognitive resources to achieve higher levels of performance.

Third, dopamine is assumed to play a role in mediating effortful behavior. Effort has been shown to be related to dopaminergic activity, specifically in the midbrain (Niv et al., 2007; Kurniawan et al., 2011; Schmidt et al., 2012; Westbrook & Braver, 2016). This is further backed up by a reduced sensitivity to incentives in Parkinson’s patients. While reward can simultaneously increase response speed and accuracy in healthy controls, the performance modulation by incentives is less pronounced in Parkinson’s patients (Manohar et al., 2015).

### Performance improvements beyond speed-accuracy tradeoffs

If effort is so hard to grasp, how then can we determine whether people engage in a task (i.e., exert effort) or not? A straightforward way to deal with this is to look at their task performance, mainly regarding response speed (e.g. reaction time, RT) and accuracy (e.g. proportion correct), and to compare it to a baseline. However, measures of speed and accuracy are tightly interrelated via a potential speed-accuracy tradeoff (SAT; Fitts, 1954; Standage et al., 2014; Heitz, 2014): For most tasks we perform, we can to some extent decide whether we emphasize speed and accept a certain number of errors, or whether we aim to respond cautiously, taking all the time we need to avoid mistakes. SAT describes this lawful relationship for a given task difficulty. Strikingly, the standard instruction in behavioral experiments to respond both as quickly and as accurately as possible allows for various degrees of freedom in this regard.

If effort indeed increases overall cognitive capacities, one might argue that it should improve performance beyond any tradeoff between speed and accuracy. Based on this more narrow conceptualization of effort, a relevant follow-up question is related to the issue of how we can determine whether a performance modulation is beyond a mere tradeoff between speed and accuracy. An intuitive and straightforward solution is to analyze the joint pattern of mean reaction times and accuracy: When an increase in accuracy goes hand in hand with an increase in reaction times, this is typically considered a speed-accuracy tradeoff (positive slope when plotting mean accuracy over mean reaction time), whereas increased accuracy combined with faster responses is typically assumed to reveal a performance improvement that clearly goes beyond any SAT (negative slope). Yet, this joint pattern of mean reaction times and mean accuracy can also be misleading (see Suppl. Fig. S1). One potential alternative are combined speed-accuracy indices (for an overview see Vandierendonck, 2017; Liesefeld & Janczyk, 2019), of which some are, however, also based on mean RT and accuracy. Another alternative are sequential sampling models, such as LATER (Carpenter & Williams, 1995; Noorani & Carpenter, 2016) or the drift-diffusion model (Ratcliff, 1978; Ratcliff et al., 2016). These approaches jointly consider reaction time distributions and error rates and help to distinguish speed-accuracy tradeoffs from other effects (Voss et al., 2004). Yet another alternative are accuracy time course analyses (often referred to as conditional accuracy functions; Heitz, 2014), which plot the probability of a correct response given a certain RT. To compare two time courses (e.g., data from two experimental conditions), responses are either sorted into quantiles or assigned to pre-defined bins. More elaborate methods allow comparing time courses without aggregating responses within in a certain time window and thus without sacrificing any information (van Leeuwen et al., 2019).

Finally, a fourth alternative to the analysis of mean RT and mean accuracy in detecting effortful behavior is the recording of eye movements as a target domain of performance. The neural network controlling saccadic eye movements contains the same midbrain structures that are also involved in motivated, goal-oriented behavior (Hikosaka et al., 2006; Hikosaka et al., 2014). Their initiation shows the same modulation with dopamine (Kori et al., 1995; Nakamura & Hikosaka, 2006), giving room for a variety of reward effects on saccadic eye movements (Hutton, 2008; Wolf & Lappe, 2021) and even revealing markers of Parkinson’s disease (MacAskill & Anderson, 2016). Importantly, eye movements are not only an alternative effector system that allows for quantifying reaction times, but the movement itself provides a manifold of relevant information for the sake of detailed performance analyses: Saccades have a unimodal velocity profile, and their peak-velocity as well as their amplitude are both sensitive to reward, allowing us to quantify motor markers of motivated, goal-directed behavior, and thus of effort (Chen et al., 2013; Kojima & Soetedjo, 2017). Further effort markers that can be derived from eye tracking technology are the saccade trajectory and pupil size. The saccade trajectory is affected by conflicting stimuli (Doyle & Walker, 2001; van der Stigchel et al., 2006) and can thus be used as an index of response conflict and the potential recruitment of cognitive control mechanisms. In this way, saccade accuracy is much more informative than merely dichotomous right/wrong responses in typical manual key press studies. Finally, the pupil size of the eye is known to be indicative of effort (e.g. Alnæs et al., 2014), linking effort and saccadic activity (Koevoet et al., 2024; Herrmann & Ryan, 2024). Taken together, eye movements provide a variety of information that allow for a detailed and comprehensive analysis of human performance and may therefore also reveal important novel insights into effortful behavior.

### Voluntary control of cognitive capacity

Contemporary research has overly focused on rewards when studying motivational processes. Yet, rewards may indirectly activate internal self-instruction, coaxing participants to invest effort. Thus, research including explicit incentives may overlook the potential sufficiency of direct instructions. Several studies have used effort instructions to investigate this possibility to mobilize capacity (Kleinsorge, 2001; Falkenstein et al., 2003; Steinborn et al., 2017; Unsworth et al 2022; Strayer et al., 2023). Kleinsorge (2001) used an instructional cue that appeared in 20% of trials and that indicated participants to speed up their responses without sacrificing accuracy. The impending response in these effort trials came along with decreased mean reaction times and increased error rates compared to the remaining 80% of trials (standard trials). A follow-up study (Falkenstein et al., 2003) replicated this behavioral pattern and showed that effort and standard trials additionally differed in event-related potentials following the instructional cue. Yet, given that participants in both studies were rewarded for speeded responses, it is not clear whether effort instructions alone were sufficient to mobilize capacities for speeded action, or whether this requires the additional presence of performance-contingent reward. Neither does any of the two studies allow for unequivocally inferring performance modulations beyond a speed-accuracy tradeoff. Building up on this work, Steinborn et al. (2017) made use of the same two trial types (effort and standard trials) without any (explicit) reward. By focusing on RT distributions rather than mean RT, this study showed that effort instructions did not speed up all responses to the same extent, but that it mainly reduced longer reaction times, thereby increasing response consistency. Again, conditions with faster responses were accompanied by a higher mean error rate. However, to foster our theoretical understanding of effort, more fine-grained performance parameters are required. As noted above, this gap may be filled by utilizing eye movement recordings.

While effects of reward on oculomotor control have been studied extensively (for reviews see Hikosaka, 2007; Madelain et al., 2011; Wolf & Lappe, 2021), and may be assumed to impact on effort mobilization, hardly any studies have addressed whether effort-related instructions *alone* (i.e., in the absence of explicit reward) can affect basic oculomotor control. Partly, this research gap may be due to the fact that eye movements are typically considered means to some ultimate goal (e.g., visual search, scene perception, reading; see Rayner, 2009), not as a target of instruction themselves (i.e., participants may typically be instructed to read faster, but not to execute their eye movements faster and more accurately at will, see also Huestegge et al., 2019). An example for a notable exception is a study by Muhammed et al (2020), who provided participants with trial-wise feedback about their peak-velocity and tested whether people can modulate their peak-velocity when asked to either respond fast or slow. Their results revealed that voluntary control of peak-velocity is possible, and that this effect can be further modulated by the presence of reward. However, their study cannot determine whether such a voluntary modulation is possible in the absence of biofeedback mechanisms. Neither does it allow for distinguishing whether people can voluntarily decrease their saccade velocities compared to normal saccade execution, or whether a voluntary up-regulation of saccadic velocity is possible. Only the latter would potentially be indicative of the notion that mobilizing spare cognitive capacities goes along with changes in peak-velocity.

### The current study

The aim of the current study was threefold. First, using the effort operationalization of performance modulations beyond speed-accuracy tradeoffs, we here test whether instructions to “try hard” and to engage in a task are sufficient to induce effortful behavior, or whether this requires the (additional) presence of an explicit performance-contingent reward. Participants had to complete a saccadic selection task that requires to inhibit an initially displayed distractor and to instead select a subsequently appearing target. In most trials (80%), participants were instructed to look at the center of the target as quickly and as accurately as possible (standard instructions in typical experimental performance tasks). Accuracy in this task refers to the avoidance of an erroneous response (e.g., looking at the distractor). Crucially, across the experiment, we interleaved sets of ten successive effort (“try harder”) trials, in which participants were instructed to mobilize all their resources and to respond even faster and make even fewer mistakes (Kleinsorge, 2001; Falkenstein et al., 2003; Steinborn et al., 2017; Unsworth et al., 2022). Each participant completed two blocks containing effort trials and standard trials, with the only difference being that in one of the two blocks correct effort trials yielded monetary reward, depending on the reaction time in that particular trial. We reasoned that telling participants to give their best and providing them with an *additional* performance-contingent reward will likely raise their performance close to the individual maximum and allows us to test whether instruction-based effort alone yields maximal effects already. Therefore, the rewarded effort condition can be considered an upper limit of performance. Additionally, we recorded a third block that only contained standard trials. This block served as a baseline and thus as a lower limit of performance (even though in the baseline trials participants are still, notably, generally instructed to respond fast and accurately, see above), thus spanning the space of potential performance improvements. Moreover, the baseline additionally serves to assure that effort trials indeed go along with an improvement in performance, and that any differences between effort and standard trials from the same block do not simply arise because participants tend to “rest” to some extent in-between sets of consecutive effort trials (i.e., a relaxation of performance in standard trials instead of a performance boost in “try harder” trials). If the latter was the case, then we should observe a difference between standard trials from the baseline and those standard trials that were interleaved with effort trials.

The second aim of the current study was to assess the potential of maintenance of effort over time. Specifically, we looked at how successful participants can maintain effort over short durations (i.e., across a set of 10 successive effort trials) as well as over longer durations (i.e., over the course of the experiment). Again, we hypothesized that maintaining effort is most successful when effort is rewarded. The third and final aim of the study was to see whether similar effects that can be found in the classic cognitive control markers of performance (i.e., proportion correct and saccadic RTs) can also be found in the dynamics of the eye movement itself, that is, in oculomotor kinematics (with respect to saccade amplitude (gain) and peak-velocity), which are well-known to respond to reward manipulations (Wolf & Lappe, 2021).

## Methods

### Participants

We recorded data of 36 participants (median age = 21 years, age range: 19–30, 7 males, 29 females). The sample size was identical to Wolf and Lappe (2023). Participants were undergraduate students from the University of Münster and were compensated with either course credit or 8€/h for participation. Participants received an additional performance-dependent reward of up to 8€ in the reward block (up to 0.10€/trial). Obtained rewards were rounded up to the first decimal after the comma and ranged from 2.10 to 4.50€ (median: 3.70€). Written informed consent was provided before testing. Participants provided written informed consent to participate in the study and regarding publishing their data. A previous version of the experiment was approved by the ethics committee of the Department of Psychology and Sport Sciences of the University of Münster. The study was performed in accordance with the ethical standards as laid down in the 1964 Declaration of Helsinki.

### Setup

Stimuli were presented on an Eizo FlexScan 22-inch CRT monitor (Eizo, Hakusan, Japan) with a resolution of 1152 × 870 pixels, an effective display size of 40.7 × 30.5 cm and a refresh rate of 75 Hz. A chin–forehead rest restricted head movements and assured a viewing distance of 67 cm. Stimulus presentation was controlled via the Psychtoolbox (Brainard, 1997; Kleiner et al., 2007) in MATLAB (The MathWorks, Natick, MA). Eye position of the right eye was recorded at 1000 Hz using the EyeLink 1000 (SR Research, Mississauga, ON, Canada) and the EyeLink Toolbox (Cornelissen et al., 2002). All stimuli were presented on a black background. The EyeLink was calibrated at the beginning of each block using a 9-point calibration protocol. A drift correction was applied every 100th trial if the average gaze deviation from the fixation cross was above 2.5 deg during the initial fixation period.

### Procedure, stimuli and design

Trial procedure (Fig. 1A) was mostly equivalent to Wolf and Lappe (2023): Each trial started with an initial text displayed at the screen center for 1 s, indicating whether the upcoming trial will be an effort trial (green font, “anstrengen”, which translates to “try hard” or “make an effort”) or a standard trial (red font, “standard”). Subsequently, four dark blue discs appeared, together with a white fixation cross at the screen center. The fixation cross was a combination of hair cross and bull’s eye (Thaler et al., 2013). The four discs had a radius of 2 deg, were arranged in a square and each disc center was 11.3 deg away from the screen center. One of the discs turned white after 720 to 800 ms. After additional 120 ms, one of the two adjacent discs turned gray. This was the target. Participants were instructed to look at the center of the target disc. A disc was considered selected if horizontal and vertical gaze were each less than 2 deg away from the disc center. Trials were labelled correct if the first disc selected was the target disc and if this selection occurred within 1250 ms. Trials in which any other disc was selected were labelled as errors. If no disc was selected within 1250 ms after target onset, the trial was labelled as too slow, in which case “too slow” (feedback) was shown at the end of a trial.

**Figure 1.**
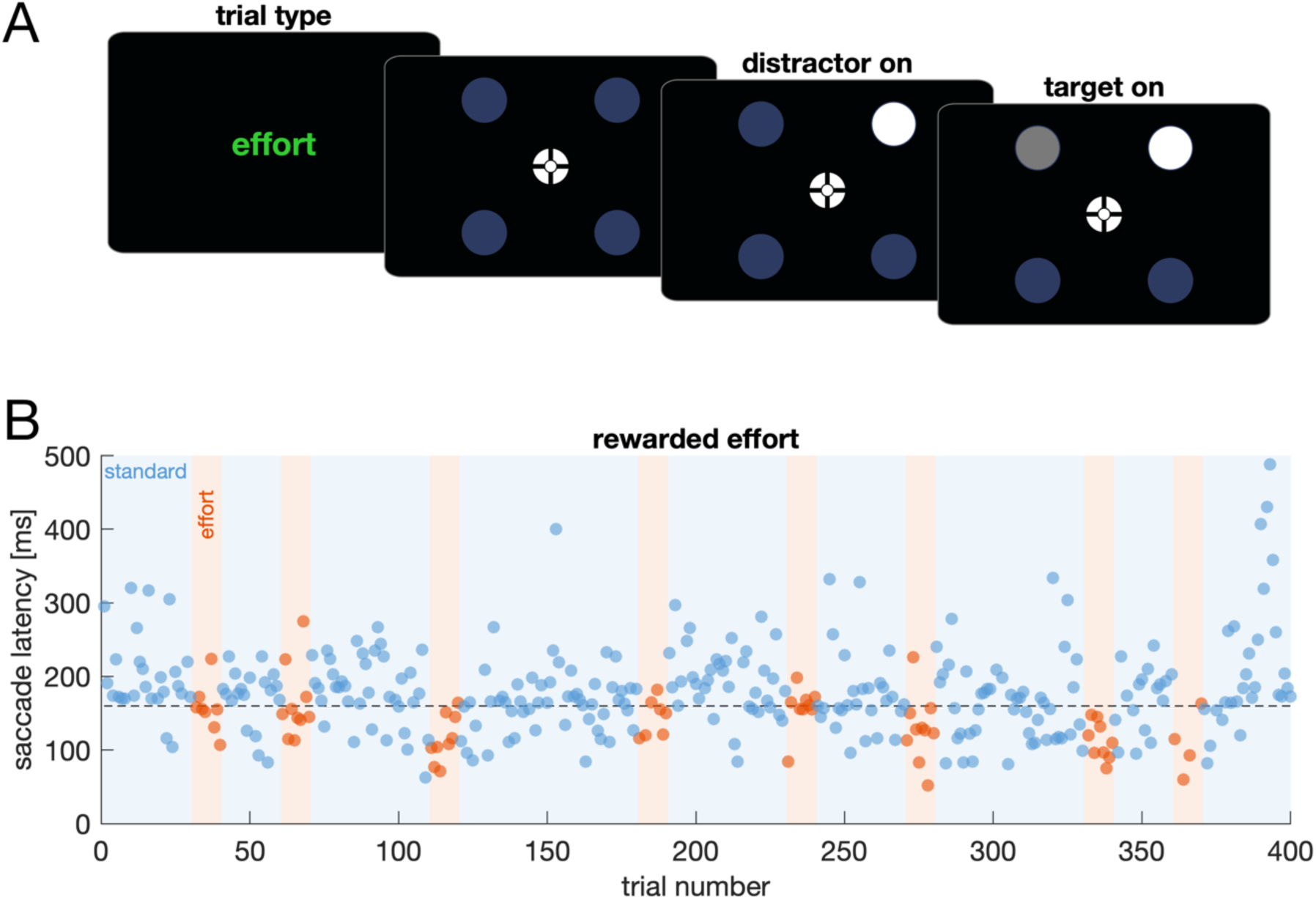
Trial procedure and individual data. (**A**). A text at the beginning of each trial indicated whether the upcoming trial was a standard trial or an effort trial. Afterwards, a fixation cross and four discs appeared. The disc turning white indicates the distractor. The disc turning gray indicates the target. (**B**) Saccade latencies over the duration of one block (rewarded effort) for one exemplary participant. Each dot denotes the data of one trial. Data from erroneous responses are missing. Dot color as well background shades denote standard trials (blue) and effort trials (orange). The horizontally dotted line indicates the mean latency from the baseline block.

The experiment comprised three blocks (baseline, unrewarded effort, rewarded effort). The baseline block only consisted of standard trials, while the other two blocks each consisted of both standard and effort trials. Each block contained 400 trials and was recorded on a different day to minimize carry-over effects on performance without resorting to a potentially underpowered group design. The block order was balanced across participants. The two effort blocks contained two different trial types: standard trials (320 trials) and effort trials (80 trials). In standard trials, participants were instructed to respond as quickly and as accurately as possible and told that this implies responding as quickly as possible without making mistakes. In effort trials, participants were instructed to “give everything” to be even quicker and to make even fewer mistakes (and thus to “*give 110%*”).

Effort trials were always presented in sets of ten successive trials (Fig. 1B). For every 50 trials, effort trials were randomly positioned at one of three possible positions (trials 11–20, 21–30 or 31–40). Therefore, participants completed 20 to 60 standard trials in-between two sets of effort trials.

In the rewarded effort block, participants received a performance-contingent reward for correct effort trials, in line with established procedures to study reward on oculomotor control (see Manohar et al., 2015; Wolf & Lappe, 2023). The maximal reward per trial was 0.1€ and decayed with increasing reaction time. The strength of the decay depended on the mean latency of the preceding eight trials to assure a comparable difficulty across individuals as well as across the course of the experiment (Wolf & Lappe, 2023). Feedback was provided 0.8 deg above the target disc as soon as that disc was selected.

### Data analysis

Saccade onsets and offsets were defined using the standard EyeLink 1000 algorithm. For the analysis of saccade latency, amplitude, and peak-velocity, we only considered correct responses with amplitudes between 5–20 deg, and peak-velocities ranging from 100–900 deg/s. We computed saccade gain by dividing saccade amplitude by target eccentricity (11.3 deg). Hence, a saccade gain of 1 indicates that saccade amplitude coincides with target eccentricity. Saccade velocity was obtained by differentiating gaze position. The peak-velocity of a saccade is strongly determined by its amplitude (Bahill et al., 1975; Gibaldi & Sabatini, 2021). To reveal peak-velocity modulations beyond changes in saccade amplitude, we partialled out the effect of amplitude: For every individual we regressed peak-velocities on saccade amplitudes and computed the residual peak-velocity (Muhammed et al., 2020). As a proof of concept, we computed the correlation between residual peak-velocity and saccade amplitude for every individual. The resulting median correlation was *r* = 0.004 and thus close to zero. All statistical analyses are based on residual peak-velocity, except for the main sequence analysis (see below) which was based on absolute peak-velocity values.

Saccade latency, saccade gain, and residual peak-velocity were compared using a 2 x 2 repeated-measures ANOVA with the two factors effort instruction (effort versus standard) and reward (rewarded versus unrewarded). That is, effort and standard trials were compared in both the unrewarded effort block and the rewarded effort block. Additionally, conditions were compared against baseline (i.e., standard trials in the baseline block) using paired sample t-tests or Wilcoxon signed rank tests if normality was violated.

To reveal whether effort instructions sped up the whole reaction time distribution or whether they selectively affected long-latency responses (Steinborn et al., 2017), we computed delta plots. Therefore, we computed a cumulative distribution function (CDF) using 39 quantiles for each experimental condition. A delta plot is constructed by comparing two CDFs (in our case standard trials versus effort trials for each respective block). Delta plots display the difference between two corresponding quantiles over the mean of the same two quantiles. An overall speed-up would result in a delta plot with a slope around zero, whereas a selective speed up of long-latency responses would result in a positive slope. We applied the logic of delta plots to the analysis of saccade latencies and peak-velocities.

Accuracy time courses were compared using the SMART procedure (smoothing method for the analysis of response time courses; van Leeuwen et al., 2019): First, the data of each individual is temporally smoothed. Second, a weighted time course is determined that considers the individual data distribution. Third, two time courses are compared using cluster-permutation testing (Maris & Oostenveld, 2007). Time courses were compared in a time window between 0 and 300 ms after target onset, using a Gaussian kernel of 16 ms and 1000 permutations per test. The same analysis was applied to the saccade main sequence, thus analyzing peak-velocity as a function of saccade amplitude. Here, we compared peak-velocities in an amplitude range from 7 to 13 deg, with a Gaussian kernel of 0.48 deg and 1000 permutations per test.

To analyze the maintenance of effort over time, we analyzed oculomotor markers (saccade latency, peak-velocity, saccade gain) over time using linear mixed models (LMMs). To analyze effort maintenance within the set of 10 successive effort trials, we computed LMMs for effort trials, with fixed effects of reward (rewarded versus unrewarded) and trial number within effort set (1 to 10) as well as random intercepts for participant. A separate LMM was computed for saccade latency, peak-velocity and saccade gain. We were interested in the main effects of time, and the interaction of time and reward. We report slopes (i.e. change over time) together with their confidence interval. The slope is given by the fixed effects estimates (main effect of time) as well as trends (conditional upon individual reward condition). To analyze the maintenance of effort over the course of the experiment, we again used LMMs with the fixed effect variables reward (rewarded vs unrewarded), effort (effort vs standard) and trial number (1 to 400), with random intercepts for participants. Again, we only report main effects of time (i.e. trial number) and its interactions. The LMMs were run using JASP (Version 0.18.3).

## Results

### Effort instructions speed up responses

Figure 2A shows violin plots for saccade latencies. Responses were initiated earlier when instructions emphasized effort, *F*(1,35) = 89.72, *p* < 0.001, 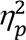 = 0.719. The effect of effort instructions was strongest when reward was present, *F*(1,35) = 13.47, *p* < 0.001, 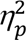 = 0.279. Effort instructions decreased response latencies by 10 ms (equivalent to 6%) in the unrewarded case, *t*(35) = 5.32, *p* < 0.001, *d* = 0.89, and by 25 ms (equivalent to 14%) in the rewarded case, *t*(35) = 7.50, *p* < 0.001, *d* = 1.25. We then addressed the issue of whether effort instructions sped up responses or whether participants slowed down responses in-between effort trials. To distinguish between these two possibilities, we compared data from the four experimental conditions with a baseline condition that contained standard trials only: Here, no differences were observed between baseline and standard trials from any of the two effort blocks, unrewarded: *t*(35) = 0.29, *p* = 0.777, *d* = 0.05, rewarded: *t*(35) = 0.64, *p* = 0.526, *d* = 0.11. In contrast, latencies in effort trials were lower than baseline, both in the unrewarded, *t*(35) = 2.17, *p* = 0.037, *d* = 0.36, and the rewarded case, *t*(35) = 4.90, *p* < 0.001, *d* = 0.816, suggesting that effort instructions did indeed speed up responses.

**Figure 2.**
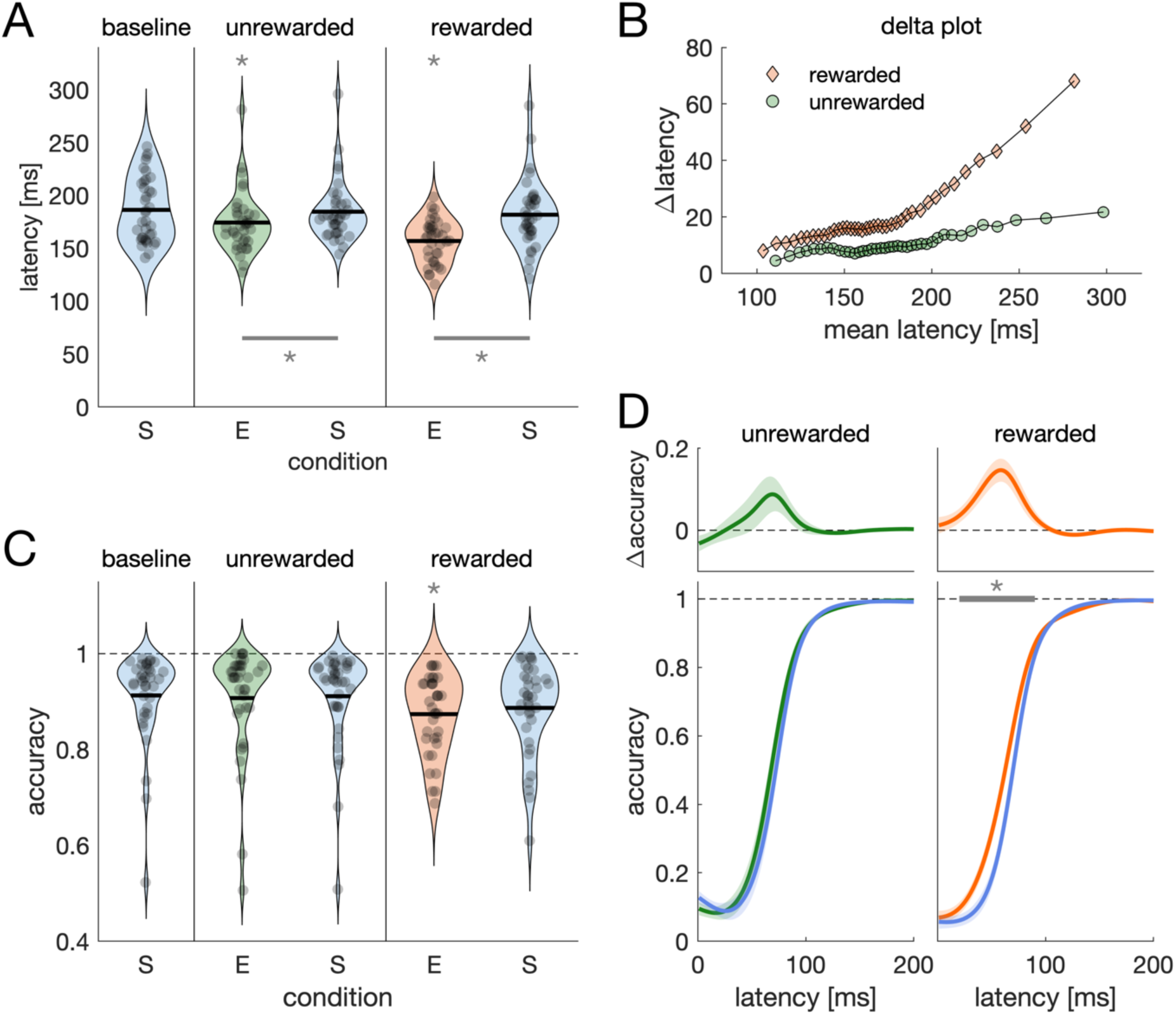
Cognitive markers of speed and accuracy. **(A)**. Violin plot of saccade latencies for the three blocks (baseline, unrewarded effort, rewarded effort) and the two trial types (E: effort trials; S = standard trials). Each dot denotes the mean of one individual, black horizontal lines indicate the aggregated means. Asterisks above a condition denote a significant difference from baseline, asterisks with lines below conditions denote a significant difference between trial types within one block. **(B)** Delta plot for the latency difference in cumulative probability between effort and standard trials. **(C)** Violin plot for mean accuracy (i.e. proportion correct responses). Same conventions as in (A). **(D)** Accuracy time courses in the unrewarded (left) and rewarded effort block (right). Shaded regions are 95% confidence intervals that result from comparing the depicted time courses (van Leeuwen et al., 2019). The upper panels show the difference in between the two time courses depicted below.

To reveal whether effort instructions sped up all responses by a similar amount, or whether effort instructions selectively reduced the likelihood of long-latency responses, we conducted a delta plot analysis. Cumulative distribution functions can be found in the supplementary material (Suppl. Fig. S2). If the whole reaction time distribution was shifted, this would result in a delta plot with a close to flat slope. In contrast, if the likelihood of long-latency responses was reduced, the latency difference should be most pronounced for the highest quantiles, resulting in a delta plot with a positive slope. Delta plots can be found in Fig 2B. A positive slope can be found in both conditions. The best fitting line through the data points has a slope of 0.09 (unrewarded) and 0.29 (rewarded), respectively.

### Only purposeful effort shifts accuracy time courses

Mean accuracy values of individuals can be found in Fig 2C. Mean accuracy in rewarded effort trials was, overall, below baseline accuracy, *z* = 2.55, *p* = 0.011. Thus, judging from the mean reaction time and mean accuracy values, performance in rewarded effort trials at first sight appears consistent with a classic speed-accuracy tradeoff account: faster reactions at the expense of accuracy. However, as noted above (Suppl. Fig. S1), mean values can be misleading when judging whether a performance modulation lies within or beyond SAT. To address this issue, we computed accuracy time courses. Whereas time courses between effort and standard trials did not differ in the unrewarded context, *t* = 30.56, *t_crit_* = 133.38, time window = 68–81 ms, *p* = 0.557, accuracy time courses between effort and standard trials differed when performance-contingent reward was present, *t* = 271.54, *t_crit_* = 184.37, time window = 21–91 ms, *p* = 0.006. Consistent with these findings as well as previous reports, performance modulations beyond SAT coincide with higher rates of information accumulation as indexed by drift-diffusion modelling (Suppl. Fig. S4; Manohar et al., 2015; Wolf & Lappe, 2023), with speed-accuracy indices (Suppl. Fig. S3; Vandierendonck, 2017; Liesefeld & Janczyk, 2019) as well as with self-reports measures using the Dundee Stress State Questionnaire (DSSQ; Matthews et al., 2002).

### Effort is reflected in oculomotor kinematics

We hypothesized that effort should be reflected in reaction time distributions and accuracy time courses, but also in the kinematics of the eye movement itself. Typically, saccades do not move in a straight line from the initial fixation to the target, but systematically deviate from this direct path based upon the location of the distractor (Suppl. Figs. S5-S6), highlighting the response conflict between looking at the distractor versus looking at the target. Moreover, we expected to find a similar pattern in oculomotor markers of accuracy and speed (i.e., saccade amplitude and peak-velocity) that we did find using the classical cognitive control markers of effort (proportion correct and reaction time, see above). Thus, we expected a strong difference between effort and standard trials in the rewarded effort block, and a reduced effect in the unrewarded effort condition. The mean amplitude across all trials amounted to 10.8 deg, and the mean peak-velocity was 458 deg/s and thus in a range that would be expected from saccades with amplitudes mostly ranging from 8 to 12 deg (Gibaldi & Sabatini, 2021).

We computed saccade gains by dividing saccade amplitude by the target eccentricity (11.3 deg). The average saccade gain amounted to 0.96. Figure 3C shows violin plots of saccade gains. Gains were higher when effort was instructed, *F*(1,35) = 17.69, *p* < 0.001, 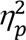 = 0.335, and, again, this difference was largest when reward was present, *F*(1,35) = 4.24, *p* = 0.047, 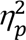 = 0.108. Saccade gains in the unrewarded block did not differ from baseline, effort trials: *t*(35) = 0.11, *p* = 0.914, *d* = 0.02, standard trials: *t*(35) = 1.29, *p* = 0.206, *d* = 0.215. However, in the rewarded block, gains from both trial types were above baseline, effort trials: *t*(35) = 5.91, *p* < 0.001, *d* = 0.985, standard trials: *t*(35) = 5.14, *p* < 0.001, *d* = 0.857.

**Figure 3.**
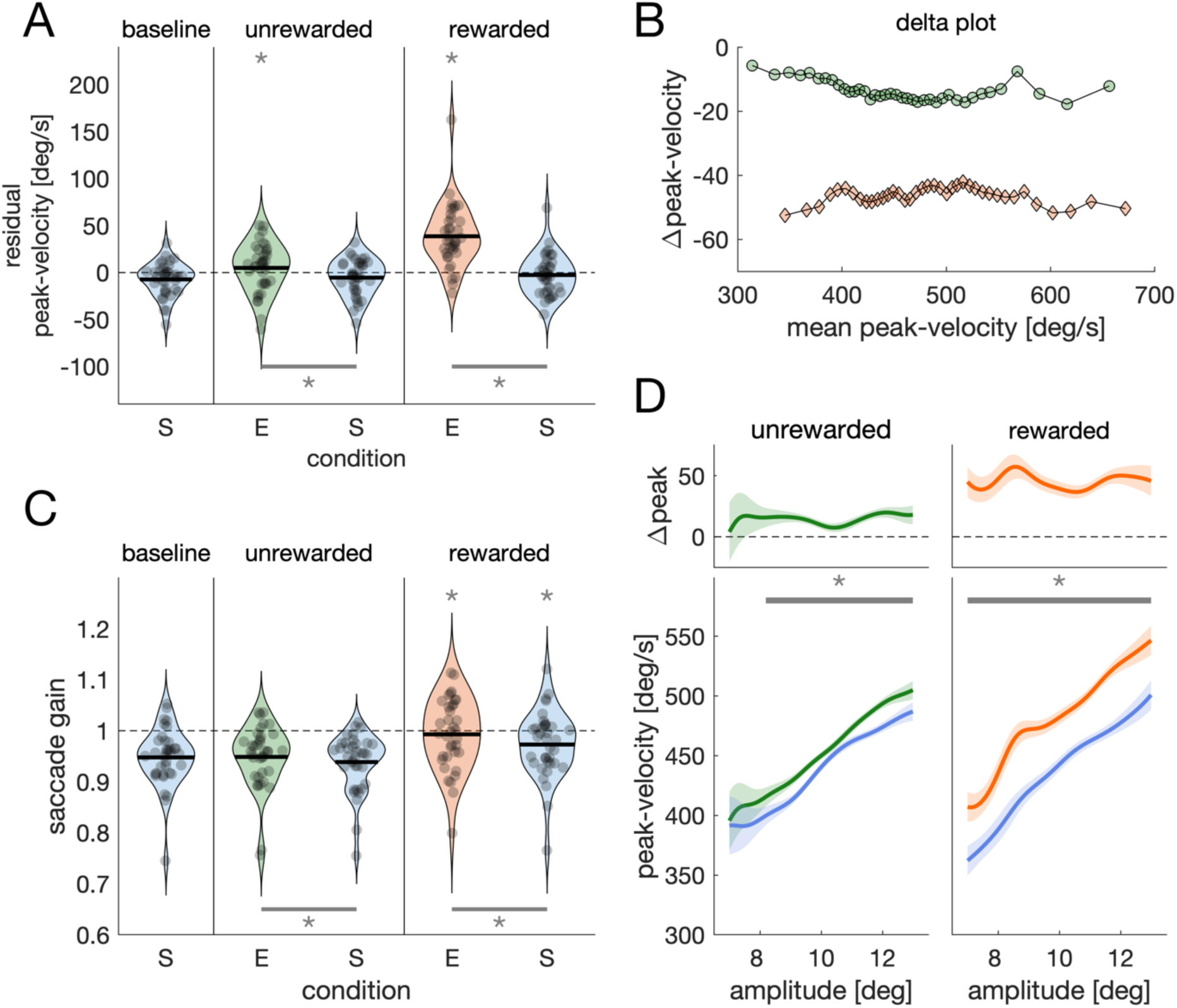
Oculomotor markers of speed and accuracy. **(A)**. Violin plot of residual peak-velocities. **(B)** Delta plot for the peak-velocity difference in cumulative probability between effort and standard trials. **(C)** Violin plot of saccade gain. **(D)** Main sequence in the unrewarded (left) and rewarded effort block (right). We again conducted a delta plot analysis to reveal whether effort instructions shifted the whole peak-velocity distribution or whether they selectively reduced data points at one end of the distribution. Delta plots can be found in Figure 3B. Here, delta plots display a slope close to zero, both in the unrewarded (−0.02) and the rewarded conditions (−0.001), suggesting a shift of the whole peak-velocity distribution.

To factor out the effect of saccade amplitude on peak-velocity (see Methods), we computed residual peak-velocities. Figure 3A shows a violin plot of residual peak-velocities. Interestingly, effort instructions coincided with increased peak-velocities, *F*(1, 35) = 8.19, *p* < 0.001, 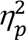 = 0.62. This effect was larger when effort was rewarded compared to unrewarded, *F*(1,35) = 44.69, *p* < 0.001, 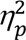 = 0.561. We compared residual peak-velocities with baseline, to reveal whether effort instructions did speed up saccade velocities or whether saccade velocities were indeed reduced in standard trials that occurred in-between effort trials. We observed no difference between baseline and standard trials from any of the two effort blocks, unrewarded: *t*(35) = 0.40, *p* = 0.689, *d* = 0.07, rewarded: *t*(35) = 0.82, *p* = 0.42, *d* = 0.14. In contrast, peak-velocities were above baseline when effort was instructed, unrewarded: *t*(35) = 2.05, *p* = 0.048, *d* = 0.34, rewarded: *t*(35) = 6.15, *p* < 0.001, *d* = 1.02. The difference in residual peak-velocity between effort and standard trials was 10 deg/s in the unrewarded case and 41 deg/s in the rewarded case.

The relationship between speed and accuracy of oculomotor kinematics (i.e., between peak-velocity and saccade amplitude) is given by the main sequence (Bahill et al., 1975, Gibaldi & Sabatini, 2021). We compared main sequences in the amplitude range that we obtained from the natural variability in saccade amplitude (Fig. 3D). Main sequences differed, both in the unrewarded, *t* = 1679.66, *t_crit_* = 425.69, amplitude range: 8.19–13 deg, *p* < 0.001, as well as in the rewarded case, *t* = 3767.95, *t_crit_* = 436.69, amplitude range: 7–13 deg, *p* < 0.001.

### Maintaining effort across short- and long-ranging timescales

We hypothesized that maintaining effort was more successful when incentives were present. At first, we were interested in the maintenance of effort on a short-range timescale, namely during the ten successive effort trials. Therefore, we compared data of saccade latencies, residual peak-velocities and saccade amplitudes from rewarded and unrewarded effort trials over time using linear mixed models (LMMs). The data are plotted in Figure 4A-C. We were exclusively interested in the main effect of time, and the interaction between time and reward. Whereas the reduction in saccade latencies relative to standard trials was relatively constant over time when reward was present, β = −0.10 ms per trial [-0.78 0.59], latency reduction decayed without reward, β = 1.22 ms per trial [0.55 1.90], interaction time x reward, *t*(4842.6) = 2.70, *p* = 0.007. Thus, maintaining an increased response speed was more successful when reward was present. In contrast, we found no interaction between reward and time, neither for peak-velocities, *t*(4843.0) = 0.82, *p* = 0.415, nor for saccade gain, *t*(4841.6) = 0.79, *p* = 0.428, but observed a main effect of time, indicating decreasing peak-velocities, β = −1.44 deg/s per trial [−2.26 −0.61], *t*(4844.0) = 3.41, *p* < 0.001, as well as decreasing saccade gain over time, β=-0.002 [−0.0032 −0.0008] per trial, *t*(4841.9) = 2.99, *p* = 0.003.

**Figure 4.**
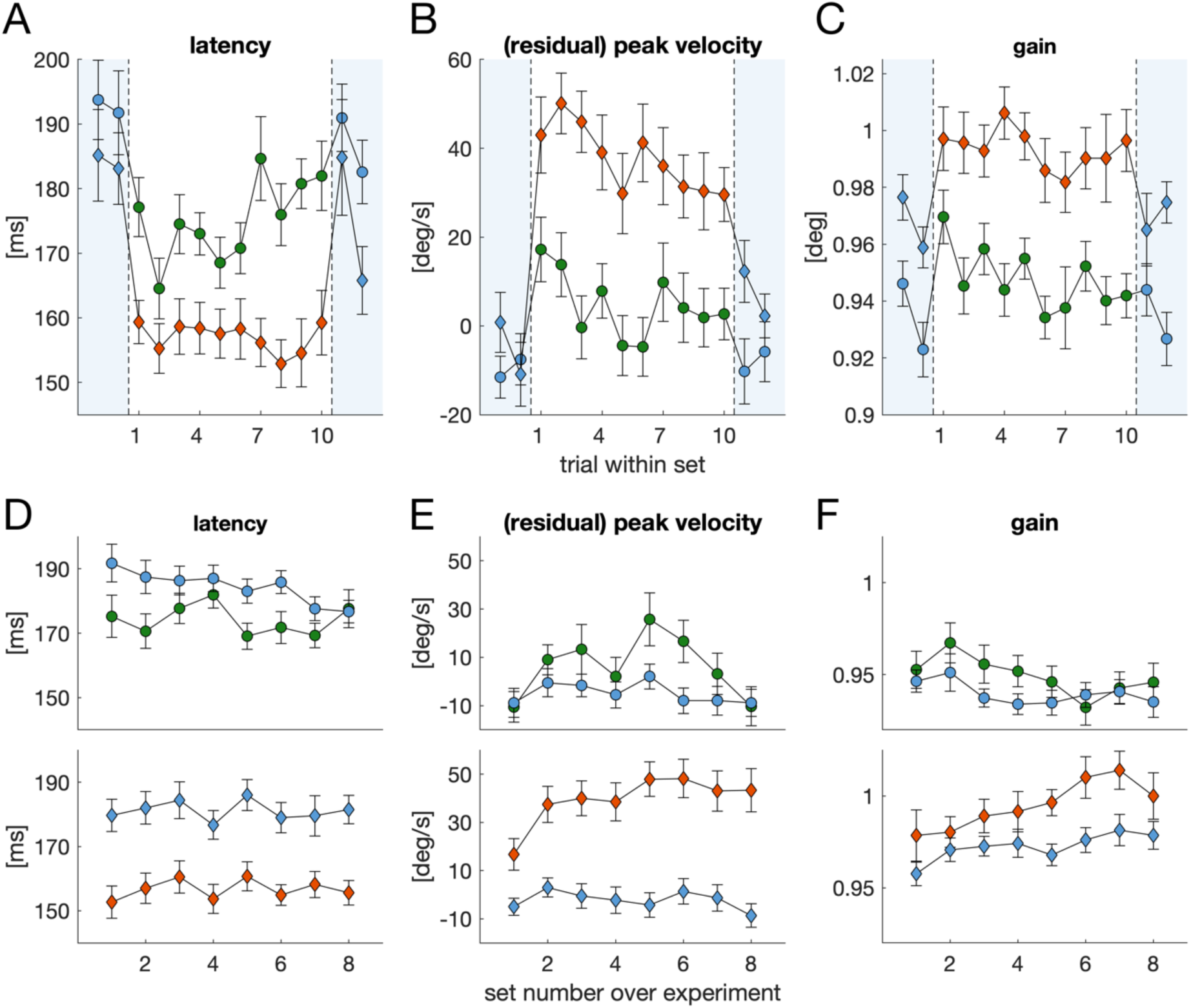
Maintaining effort across short-range and long-range timescales. (**A-C**). Modulation of oculomotor markers during sets of ten successive effort trials (blue: standard trials; green: unrewarded effort trials; orange: rewarded effort trials). (**D-F**) Modulation of oculomotor markers over the course of the experiment. Data are divided into sets, with each set containing 50 trials (10 successive effort trials and 40 standard trials). Error bars are standard errors of within-participant variability (Cosineau, 2005).

Next, we were interested in maintaining effort across long-ranging timescales, namely over the course of the 400 trials of the experimental session (time-on-task). Again, we computed LMMs, this time with the fixed effect variables reward (rewarded vs unrewarded), effort (effort vs standard) and trial number. Figure 4D-F shows mean values of effort and standard trials within each 50 trials, separately for saccade latency, peak-velocity and saccade gain. On average, saccade latencies decreased over the course of the experiment, β = −0.011 ms per trial [−0.019 −0.003], *t*(24261.4) = 2.81, *p* = 0.005. Yet, this main effect is mainly driven by standard trials in the unrewarded condition, β = −0.033 ms per trial [−0.043 −0.023]. For peak-velocities, the LMM revealed a three-way interaction between reward, effort and time, *t*(24264.6) = 2.09, *p* = 0.037. When reward was present, peak-velocities in effort trials increased over the course of the experiment, β = 0.043 deg/s per trial [0.014 0.073], whereas a tendency of a trend in the opposite direction could be observed in standard trials of the same block, β = −0.012 deg/s per trial [-0.027 0.002]. In the condition without reward, we observed no linear trend over the course of the experiment, neither for effort, β = 0.002 deg/s per trial [-0.027 0.031], nor for standard trials, β = 0.004 deg/s per trial [-0.019 0.010]. The pattern in saccade gain over time can best be summarized by an interaction between reward and time, *t*(24260.7) = 6.25, *p* < 0.001. Whereas saccade gain increased over the course of the experiment when effort trials were rewarded, both for effort trials, β = 7.66×10^-5^ per trial [3.48×10^-5^ 1.18×10^-4^], but also for standard trials, β = 6.00×10^-5^ per trial [1.06×10^-5^ 3.93×10^-5^], saccade gain decreased in the unrewarded condition, both for effort trials, β = −4.277×10^-5^ per trial [-8.40×10^-5^ −1.55×10^-6^], and standard trials, β = −2.98×10^-5^ per trial [-5.06×10^-5^ −9.06×10^-6^].

## Discussion

We tested three main research questions: First, do mere effort instructions induce effortful behavior as evidenced by response speed and accuracy, or does this additionally require the presence of performance-dependent reward? Second, how is effort maintained across short-ranging and long-ranging timescales? Third, can we find the same effort-related pattern in specific, more fine-grained oculomotor markers (saccade amplitude and peak-velocity) as in the classic cognitive markers of speed and accuracy (RT and proportion correct)?

The results were clear-cut. Overall, effort instructions decreased saccadic reaction times (Fig. 2A), replicating earlier findings in the context of manual response control (Kleinsorge, 2001; Falkenstein et al., 2003; Steinborn et al., 2017). As we utilized eye movements as a response system, this allowed us to additionally assess kinematic markers of oculomotor control. Indeed, effort instructions increased saccadic peak-velocity and spatial gain (amplitude) (Fig. 3). All these effects were strongest when reward was present. Thus, mere effort instructions appear to be sufficient to speed up responses and modulate eye movement kinematics, yet not to the same extent as with an additional performance-contingent reward (Suppl. Table T1). Furthermore, using a baseline measure allowed us to distinguish between two possibilities, namely either that participants up-regulate their performance in effort trials or that participants rest in-between sets of consecutive effort trials. The results unequivocally support the former.

When looking at mean accuracy (i.e. mean proportion correct, Fig. 2C), the combination of effort instructions and reward resulted in the lowest overall accuracy (proportion correct). Thus, based on mean accuracy alone, one might conclude that, as accuracy was worst in rewarded effort trials, performance in this condition is consistent with the assumption of a speed-accuracy tradeoff. However, when looking at the specific accuracy time courses, a different pattern emerges (Fig. 2D), revealing a shifted time course and thus a performance improvement *beyond* a mere speed-accuracy tradeoff. For unrewarded effort, we found a similar but somewhat smaller tendency towards a shifted accuracy time course, yet this difference remained at the descriptive level. Indeed, the oculomotor kinematic marker of accuracy, saccade amplitude, was more consistent with the pattern found in accuracy time courses rather than the pattern found in mean accuracy. Thus, if one considers accuracy time courses rather than mean accuracy, a consistent pattern can be found in markers of cognitive control and oculomotor markers, further highlighting that solely relying on mean accuracy analyses might be ill-advised in the context of specifying mechanisms underlying effort-based performance (Suppl. Fig. S1). The present findings therefore provide evidence for a relationship between cognitive effort and motor control, a prediction that follows from neurocomputational models of cognitive control and effort investment (Verguts et al., 2015; Silvetti et al., 2023).

Our experiment was specifically designed to reveal aspects of effort maintenance. Rather than randomly interleaving effort and standard trials, we interleaved sets of ten successive effort trials. This allowed us to test the maintenance of effort markers over the duration of this set of consecutive effort trials. For peak-velocity and amplitude, we found a general decline over the set of consecutive effort trials. Given the smaller effect of mere effort instructions without reward, values at the end of the effort set closely resembled values from subsequent standard trials, whereas in the rewarded case, an offset between effort and standard trials remained present at the end of the set (Fig. 4B-C). For saccadic reaction time, a performance decline could only be detected in the unrewarded effort condition (Fig. 4A), suggesting that reward helped to maintain the speed-up in reaction time. We also assessed the maintenance of effort over the whole duration of the experiment. This analysis revealed that only when reward was present, the difference between effort and standard trials could be maintained throughout the experiment (Fig. 4D-F).

On the aggregated level, the general cognitive indices, RT and proportion correct, and the specific oculomotor kinematics pointed into the same directions. However, a closer look also revealed some important differences between proportion correct and reaction time on the one hand and oculomotor kinematics on the other hand. For example, the delta plot analysis revealed that effort instructions particularly reduced the likelihood of long-latency responses, replicating similar findings of Steinborn et al (2017). This pattern was even more pronounced when reward was present (Fig. 3B). On the contrary, the delta plot analysis for peak-velocity revealed flat slopes, indicating that the whole peak-velocity distributions were shifted. This highlights that effort mobilization impinges differently on saccadic reaction times and peak-velocities. Whereas effort mobilization is reflected in a lower number of RT lapses and thus leaves room for the possibility that only a subset of (otherwise slow) trials is affected by effort mobilization, the peak-velocity results clearly suggest that all trials are affected by effort mobilization. Moreover, it was possible to maintain a lower RT level over the duration of a set of effort trials (Fig. 4A), whereas this was not possible for peak-velocity (Fig. 4B), potentially pointing out that difficulties in maintaining effortful behavior can be first detected in peak-velocities (Di Stasi et al., 2013) prior to affecting saccadic latencies. Taken together, this might imply that oculomotor kinematics such as peak-velocity might represent a more sensitive marker of effort mobilization and utilized capacity than reaction times. Indeed, peak-velocity outperformed reaction time and saccade amplitude in discriminating between effort and standard trials (Suppl. Fig. 7). Amplitudes on the other hand appeared to be sensitive to a global activation, given that rewarded effort showed a transfer to standard trials, resulting in higher saccade amplitudes in the rewarded effort block (Fig. 3C). Thus, our results support the view that peak-velocity measurements (corrected for amplitude) or the main sequence relationship itself represent key measurements to detect whether people engage in a task and can invest cognitive resources (Fig. 3D, Di Stasi et al., 2012; Gibaldi & Sabatini, 2021). Moreover, statistically speaking, detecting a difference in peak-velocities (or other oculomotor markers) more likely requires fewer trials than detecting a difference in accuracy (proportion correct), considering the difference in the variables’ scale (metric versus dichotomous). Taken together, these results demonstrate that the utilization of eye tracking is indeed able to uniquely advance our theoretical understanding of the mechanisms underlying effects of effort mobilization on performance.

The present findings show that eye movement behavior can differ based on self-instruction and is thus susceptible to direct voluntary control. This might have implications for studies comparing different groups of people that differ in their level of engagement, and thus in how much cognitive resources they devote to a task. Such a difference might, for example be observed when comparing athletes and non-athletes (Chen et al., 2024), potentially explaining why athletes have a lower mean saccade latency and a tendency for a higher mean error rate. Moreover, our results extend the findings by Muhammed et al. (2020) by showing that participants can up-regulate saccade peak-velocity, and that this modulation can be observed in the absence of feedback (i.e., in the unrewarded effort condition). In our study, voluntarily increasing peak-velocities was even more successful when participants received feedback about the obtained reward at the end of an effort trial. Hence, with the present experiment we cannot ultimately distinguish whether the difference between rewarded and unrewarded effort is due to reward, due to feedback or the combination of the two. However, we consider it unlikely that feedback is the driving factor causing the difference in performance. A recent study using the same task (Wolf & Lappe, 2023) compared accuracy time courses when participants received different forms of feedback about their performance. A shift in accuracy time courses was only observed when participants received feedback about obtained reward, but not when performance was fed back in terms of image quality or task difficulty (Wolf & Lappe, 2023), showing that feedback per se cannot be the reason for the difference between our rewarded and unrewarded effort condition. Instead, it emphasizes that the presence of reward is the main driving factor.

Posner (1976) described how information processing in chronometric tasks is affected by increased effort or an alerted state compared to standard conditions. When a target appears, participants begin processing the information by analyzing the stimulus and mapping it to semantic rules. One idea is that intensified focus speeds up this processing, which leads to faster information accumulation, which in turn leads to quicker reactions. This idea is consistent with information accumulation in sequential sampling models (Ratcliff et al., 2016), rendering, for example, the drift rate in the drift-diffusion model another potential marker of effort. A second possibility to speed up responses would be to lower response threshold. In the drift-diffusion model, this would manifest in a lowered boundary separation. A lowered response threshold may not only result in an earlier, but also more likely in an erroneous response, qualifying this threshold as an index of SAT (Voss et al., 2004). These two possibilities to speed up responses do not contradict each other but can go hand in hand in the context of effort or reward (Suppl. Fig. S4; Wolf & Lappe, 2023). This can explain non-monotonic relationships between effort and performance, an identified weakness for using task performance as a measurement of effort (Thomson & Oppenheimer, 2022). Thus, it is the lowered response threshold which gives rise to more errors and overwrites potential performance improvements by an increased information accumulation, resulting in an overall lowered mean accuracy in terms of proportion correct. Yet, it is the information accumulation that mainly determines the underlying relationship between reaction time and accuracy, potentially modulating performance beyond a speed-accuracy tradeoff.

Recent attempts to summarize effort research (e.g. Shenhav et al., 2017) as well as attempts to point out the various approaches, definitions, and contradictions in the study of effort (Thomson & Oppenheimer, 2022) emphasize the involvement of a limited resource or capacity. How can we identify how much of that resource is devoted to the task? Here, we operationalized effort as performance modulations beyond a speed-accuracy tradeoff and showed that effort instructions in combination with reward fulfill this criterion in an exemplary manner (faster responses and, especially, shifted accuracy time course), while the overall pattern for unrewarded effort was less pronounced (faster responses but only a tendency towards a shifted accuracy time course). Whereas this operationalization might be applicable to tasks involving reaction times and errors, it might not be true in other scenarios, for example when performing mental arithmetic, when measuring task selection (Clay et al., 2022), or when measuring giving-up (Herrmann & Ryan, 2024). Yet, our present conceptualization allowed us to come up with an objective and testable prediction for the otherwise hard-to-grasp concept of effort.

Taken together, our study showed that i) mere effort instruction led to a regulation of oculomotor performance even in the absence of explicit reward, ii) the additional presence of reward had a substantial additional impact on performance beyond a speed-accuracy tradeoff, and iii) fine-grained oculomotor parameters (peak-velocity and gain) showed a particular sensitivity as indices of effort, In particular, the latter demonstrated theoretically important aspects of effort mobilization in performance that appear to be masked when only focusing on general cognitive parameters such as mean reaction time and mean proportion correct.

