## Supplementary Material for "Effort in Oculomotor Control: Role of Instructions and Reward on Spatio-temporal Eye Movement Dynamics"

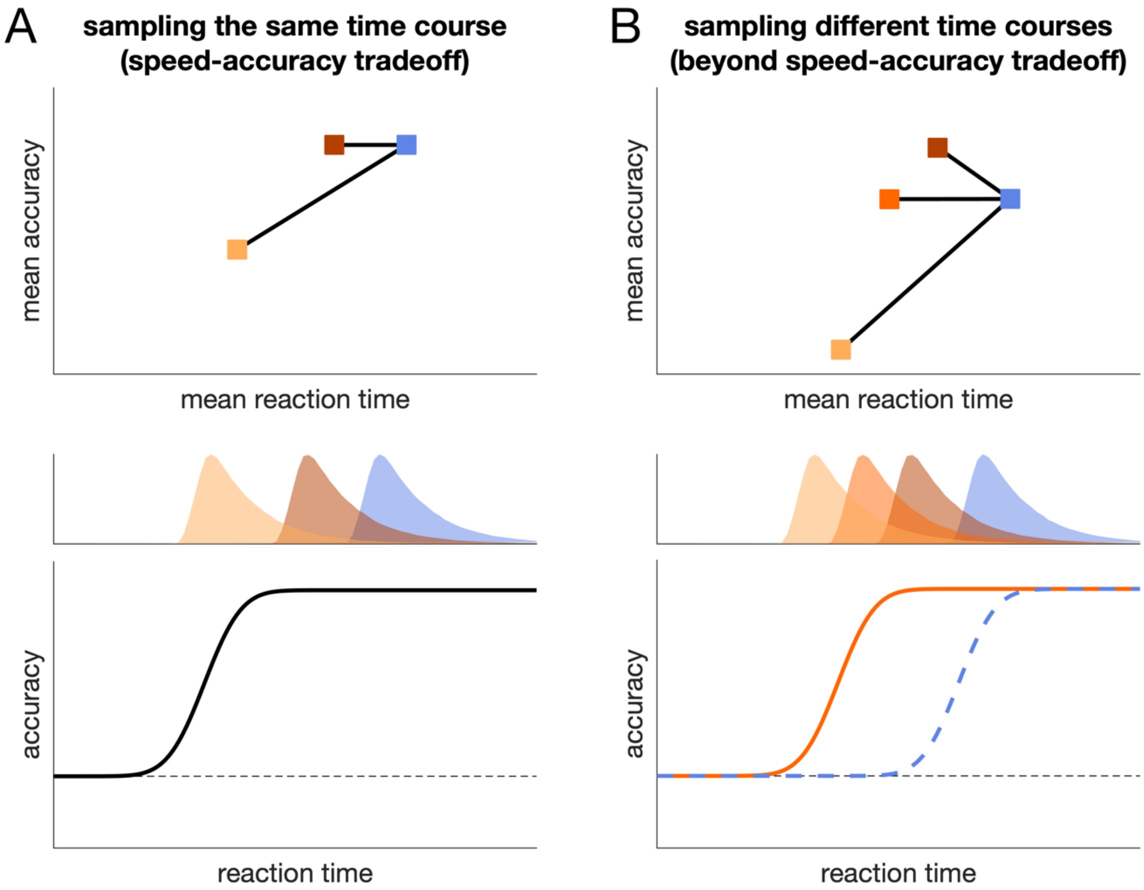


**Supplementary Figure S1. Performance modulation beyond the speed-accuracy tradeoff (SAT) is revealed by accuracy time courses – not by mean values**. (**A**). Performance modulation within SAT. When performance is reliant on the same underlying time course (lower panel), then accuracy depends on how this time course is temporally sampled (upper two panels): Compared to a baseline condition (blue), accuracy in the experimental condition (orange colors) can either decrease (i.e., SAT) or remain at the same level (i.e., ceiling effect). (**B**) Performance modulation beyond SAT. Behavior that is simultaneously faster and more accurate requires that the underlying time course is changed. Yet, even a shifted time course that reaches the same level of accuracy with faster reaction times (orange time course, lower panel) can result in better, comparable, or worse accuracy, depending on how this time course is temporally sampled. In short, a small latency increase might go along with an accuracy *increase* (a performance profile beyond a mere SAT that might otherwise go undetected), while further latency decreases might then result in accuracy decrements, yielding an overall negative correlation between latency and accuracy (when only looking at overall performance means).

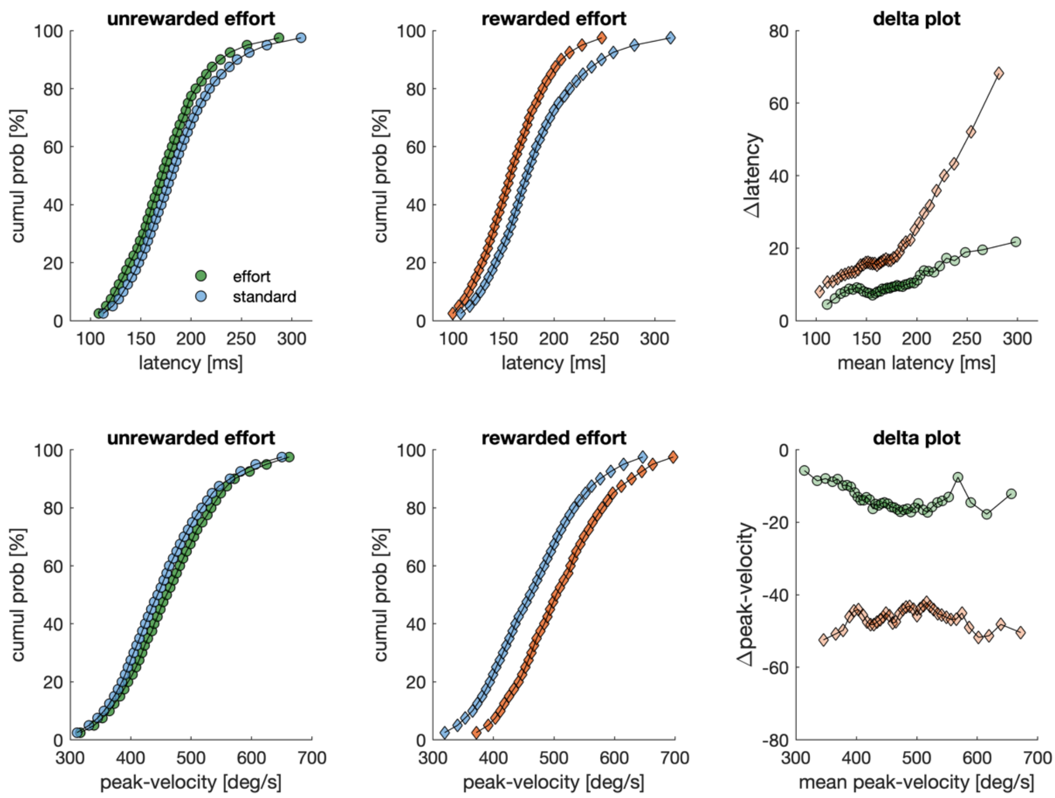


**Supplementary Figure S2. Cumulative Probability Functions**. Top row: cumulative probability plots for saccade latency (left panel: unrewarded effort; central panel: rewarded effort) and the resulting delta plots (right panel; same as Fig 3B). Bottom row: cumulative probability plots for peak-velocity (left panel: unrewarded effort; central panel: rewarded effort) and the resulting delta plots (right panel; same as Fig 4B).

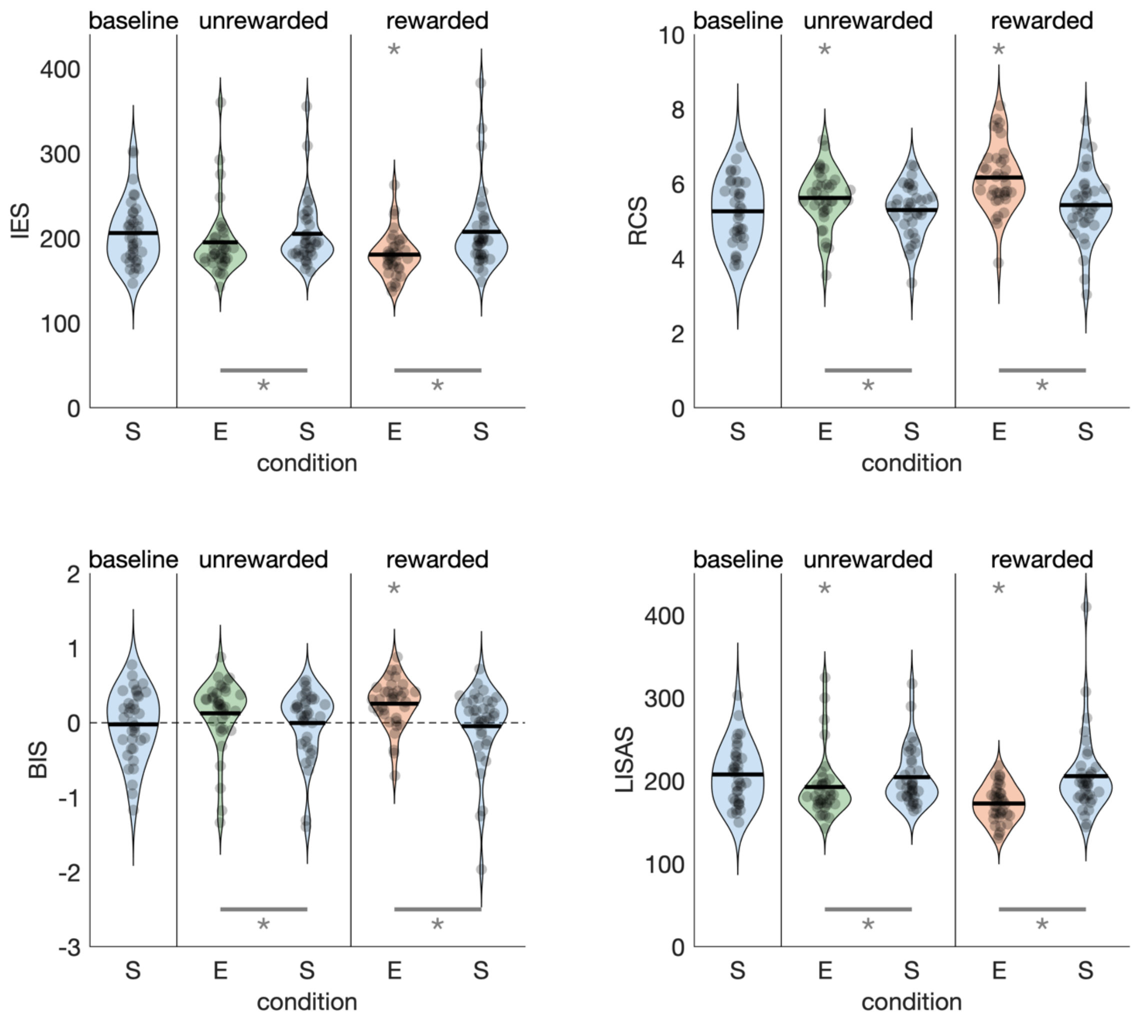


**Supplementary Figure S3. Speed-accuracy indices**. Inverse efficiency score (IES; top left), rate correct score (RCS; top right), behavioral integration score (BIS; bottom left) and linear integrated speed-accuracy score (bottom right; LISAS). Each dot denotes the score of one individual, black horizontal lines indicate the overall mean. Asterisks above a condition denote a significant difference from baseline, asterisks with lines below conditions denote a significant difference between trial types within one block (E: effort trial; S: standard trial).

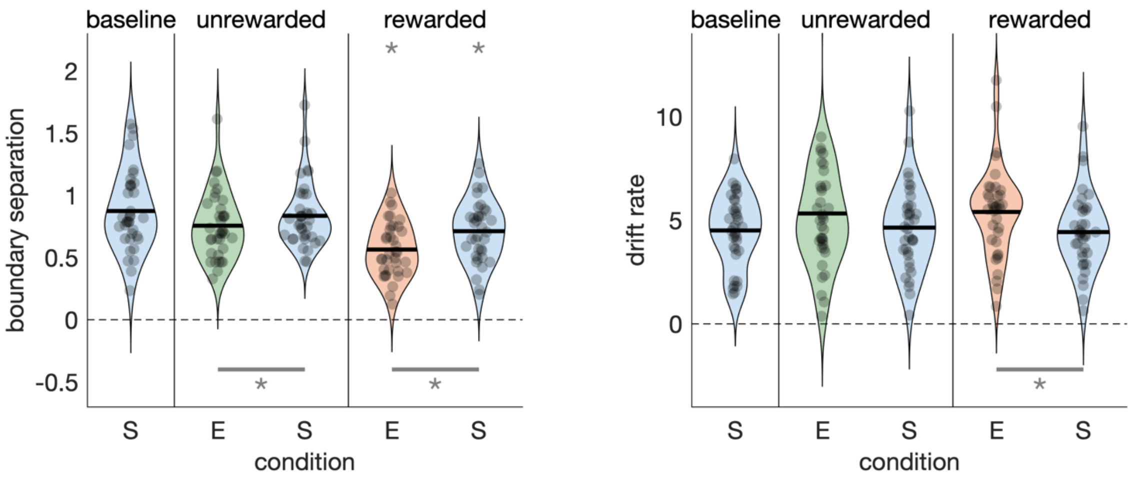


**Supplementary Figure S4. Drift-diffusion modelling results**. Violin plots for boundary separation and drift rate across the different conditions. Effort trials come along with a lower boundary separation (~ impulsive behavior / emphasizing speed) and an increased drift rate.

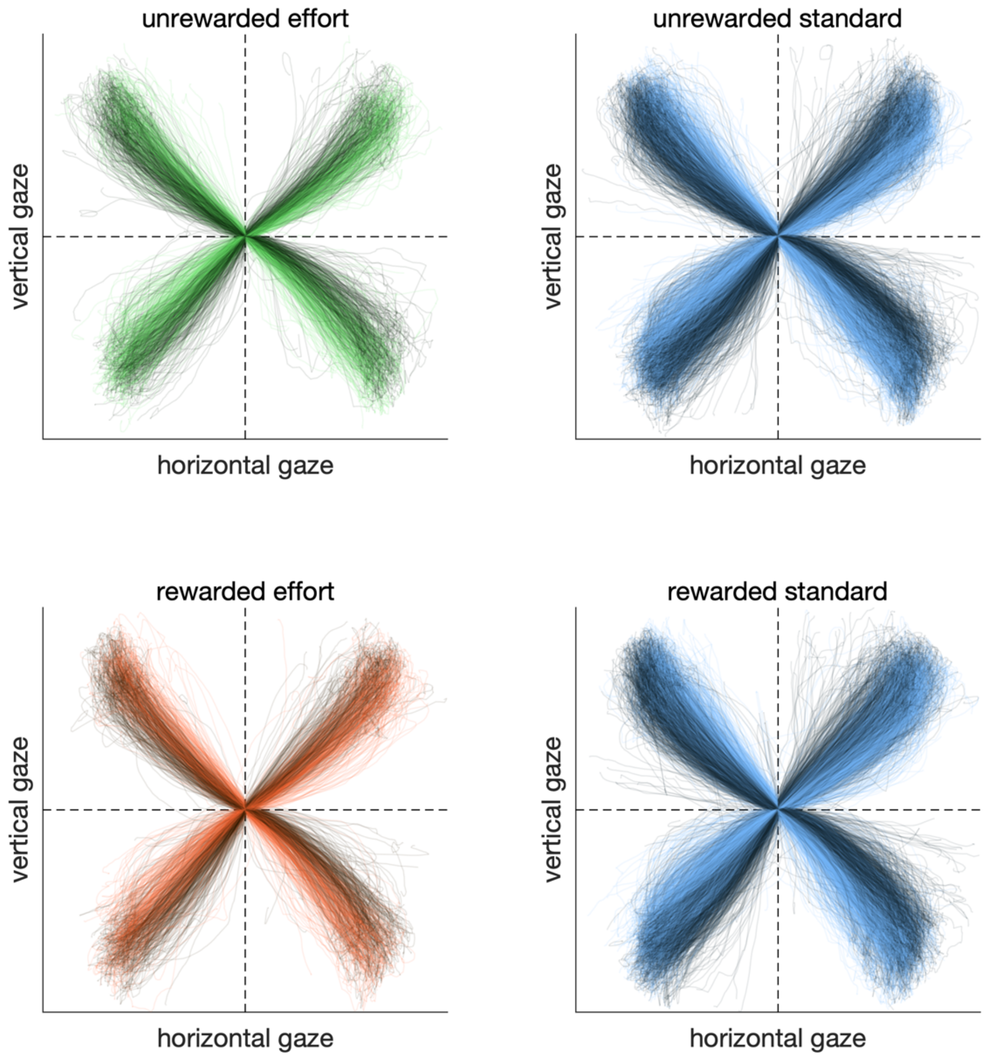


**Supplementary Figure S5. Saccade trajectories**. Saccade trajectories for effort (left panels) and standard trials (right panels) in the unrewarded (top panels) and rewarded conditions (bottom panels). Each line denotes one individual trial (correct trials only). Darker colors denote trials in which the distractor was 90 deg clockwise relative to the target location. Lighter colors indicate trials in which the distractor was 90 deg counterclockwise relative to the target location.

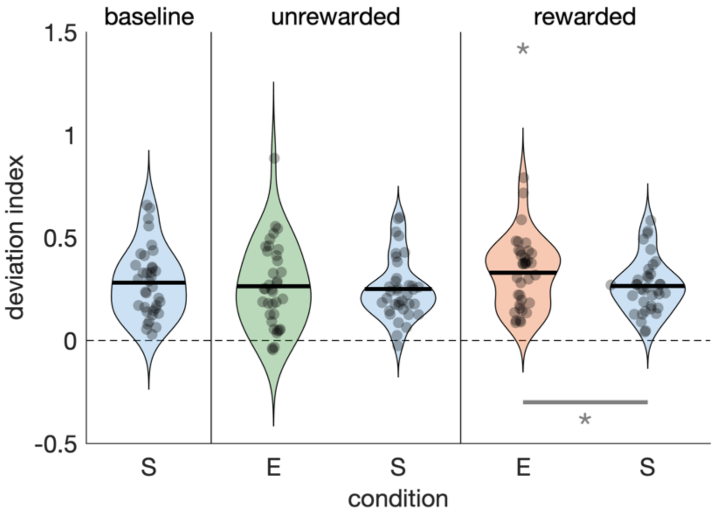


**Supplementary Figure S6. Deviation index**. Deviation indices derived from saccade trajectories (Suppl. Fig. S5). Deviation indices are computed for each trial by calculating the area in-between the saccade trajectory and a straight line between fixation cross and target. Values > 0 indicate deviation away from the distractor location.

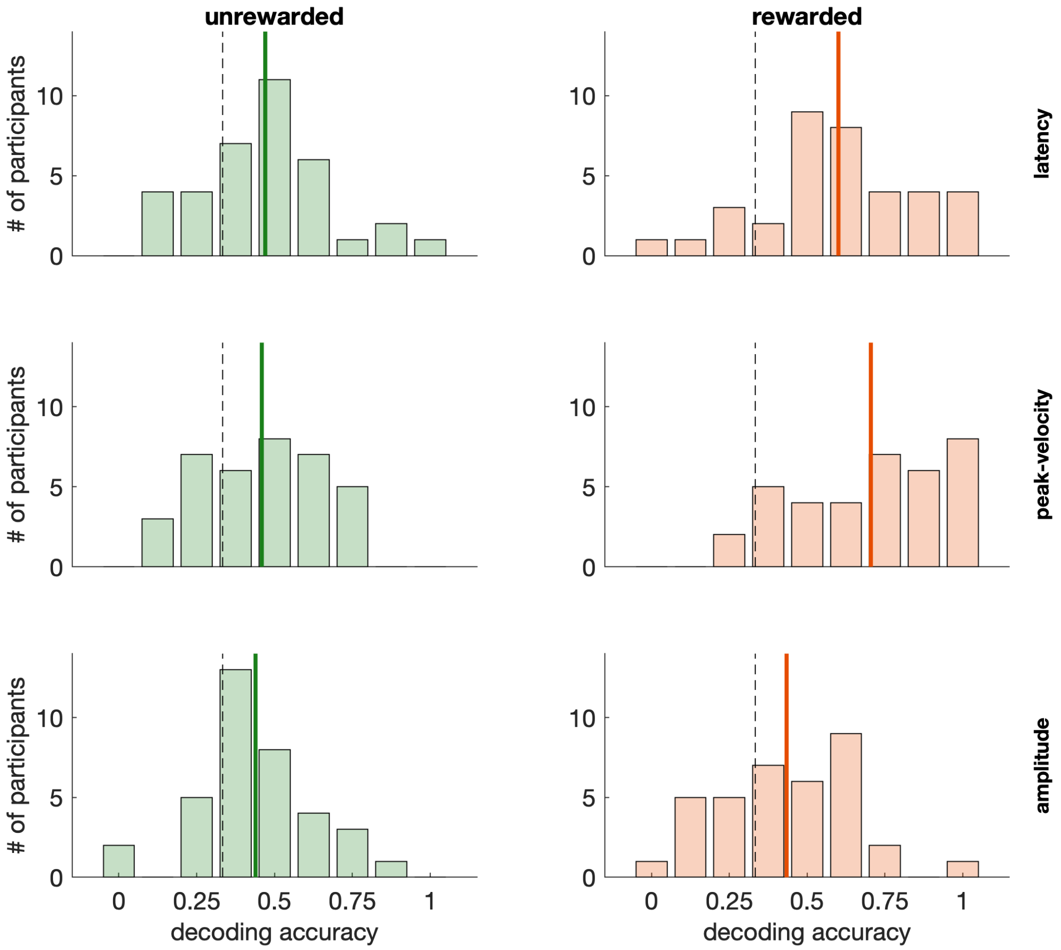


**Supplementary Figure S7. Decoding effort**. Decoding accuracy for the three oculomotor variables, separately for the unrewarded (left column) and rewarded block (right column). Dashed vertical lines indicate chance performance, solid vertical lines indicate the overall mean.

Within every 50 trials, the set of 10 consecutive effort trials could be at one of three locations (trials 11–20, 21–30, 31–40). We explored how well the three oculomotor variables can discriminate between effort and standard trials: For every participant, we computed the fraction of cases in which effort trials came along with the lowest (latency) or highest mean values (peak-velocity, amplitude) out of the three possible effort set locations (trials 11–20, 21–30, 31–40).

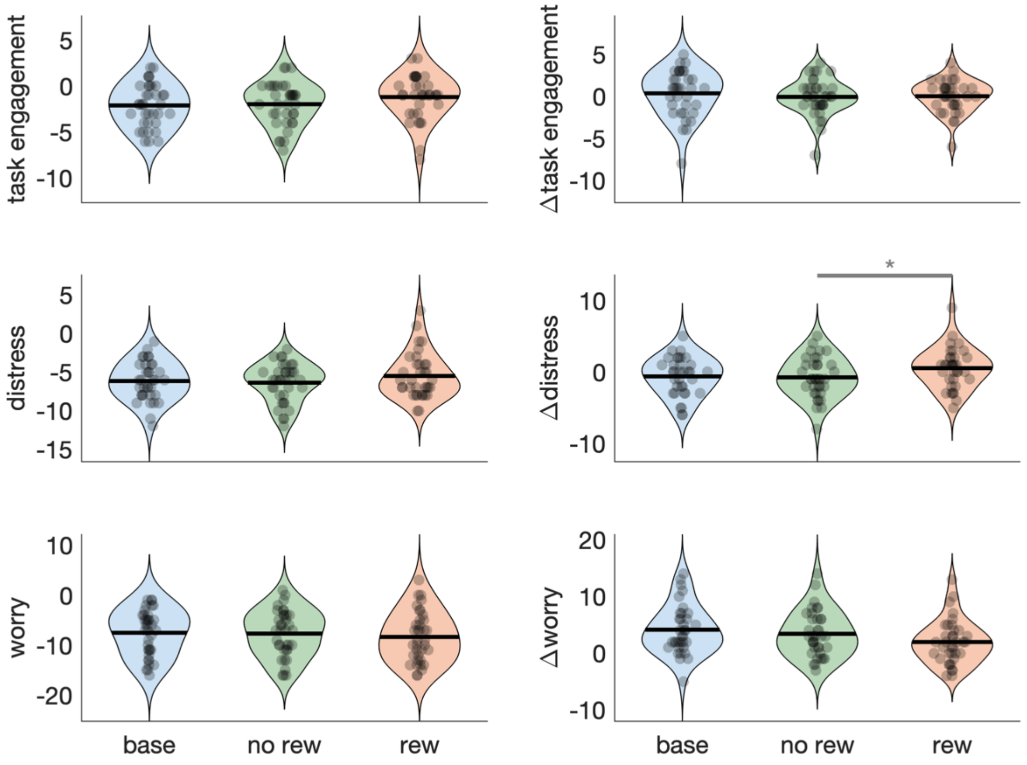


**Supplementary Figure S8. States and state changes as indexed by DSSQ**. Participants filled out the Dundee Stress State Questionnaire (DSSQ; Matthews et al., 2002) before and after each session (base: baseline; nor: unrewarded; rew: rewarded). The violin plots show the results of the post-experiment questionnaire (left column) in each of the three dimensions (task engagement, distress, worry) as well as the change relative to the pre-experiment questionnaire. The line/asterisk indicates a significant difference as indicated by a Wilcoxon signed-rank test.

Descriptively, task engagement was highest in the rewarded effort block, M = -1.2 (SD = 2.41), but neither statistically different from the unrewarded effort block, M = -1.97 (SD = 2.31), $t(34) = 1.83$, $p = 0.075$, $d = 0.18$, nor from the baseline condition, M = -2.11 (SD = 2.3), t(33) = 1.54, $p = 0.134$, $d = 0.18$.

**Supplementary Table T1. Effect size comparison between unrewarded and rewarded effort**. On average, unrewarded effort shows 60% of the rewarded effort effect size. Please note that there is no effect size metric for accuracy time courses and our main sequence analysis. Here, we used the mean relative difference as an index of effect size: For accuracy time courses, we first computed the difference between the mean time courses divided by the error at that time point. Second, we summed up all values and, third, divided by the number of time points. For the main sequence, we proceeded accordingly.

| variable | where to find data | effect size metric | effect size unrewarded effort | effect size rewarded effort | ratio |
|---|---|---|---|---|---|
| reaction time (i.e. saccade latency) | Fig. 2A | Cohen's d | 0.887 | 1.25 | 0.710 |
| accuracy time course | Fig. 2D | Area inbetween accuracy time courses | 0.363 | 1.51 | 0.240 |
| peak-velocity | Fig. 3A | Cohen's d | 0.594 | 1.359 | 0.437 |
| saccade gain | Fig. 3C | Cohen's d | 0.495 | 0.658 | 0.752 |
| main sequence | Fig. 3D | Area inbetween main sequence estimates | 5.82 | 12.11 | 0.480 |
| Inverted efficiency score | Suppl Fig. S3A | Cohen's d | 0.643 | 0.785 | 0.819 |
| rate-correct score | Suppl Fig. S3B | Cohen's d | 1.04 | 1.534 | 0.678 |
| behavioral integration score | Suppl Fig. S3C | Cohen's d | 0.648 | 0.79 | 0.820 |
| linear integrated speed-accuracy score | Suppl Fig. S3D | Cohen's d | 0.561 | 0.873 | 0.643 |
| drift rate | Suppl Fig. S4 | Cohen's d | 0.293 | 0.35 | 0.837 |
| saccadic deviation | Suppl Fig. S6 | Cohen's d | 0.089 | 0.592 | 0.150 |
